# Genomic and gene-expression comparisons among phage-resistant type-IV pilus mutants of *Pseudomonas syringae* pathovar *phaseolicola*

**DOI:** 10.1101/025106

**Authors:** Mark Sistrom, Derek Park, Heath E. O’Brien, Zheng Wang, David S. Guttman, Jeffrey P. Townsend, Paul E. Turner

**Affiliations:** School of Natural Sciences, University of California Merced, Merced, 95343, CA, USA; Department of Ecology and Evolutionary Biology, Yale University, New Haven, CT 06520, USA; Department of Cell and Systems Biology, University of Toronto, Toronto, Ontario, M5S 3B2, Canada; Centre for the Analysis of Genome Evolution & Function, University of Toronto, Ontario, M5S 3B2, Canada; Department of Biostatistics, Yale School of Public Health, New Haven, CT 06520, USA; Program in Computational Biology and Bioinformatics, Yale University, New Haven, CT 06520, USA; Program in Microbiology, Yale University, New Haven, CT 06520, USA

**Author notes:** Corresponding author’s contact details School of Natural Sciences, University of California Merced, Merced, 95343, CA, USA. Current addresses: Derek Park, Mathematical Institute, University of Oxford, Oxford, OX2 6GG, United Kingdom. Heath O’Brien, School of Biological Sciences, University of Bristol, Bristol, BS8 1TQ, United Kingdom.

**Keywords:** genomics, gene expression, *Pseudomonas syringe*, bacteriophages, ϕ6 bacteriophage

## Abstract

*Pseudomonas syringae* pv. *phaseolicola* (Pph) is a significant bacterial pathogen of agricultural crops, and phage ϕ6 and other members of the dsRNA virus family *Cystoviridae* undergo lytic (virulent) infection of Pph, using the type IV pilus as the initial site of cellular attachment. Despite the popularity of Pph/phage ϕ6 as a model system in evolutionary biology, Pph resistance to phage ϕ6 remains poorly characterized. To investigate differences between phage ϕ6 resistant Pph strains, we examined genomic and gene expression variation among three bacterial genotypes that differ in the number of type IV pili expressed per cell: ordinary (wild-type), non-piliated, and super-piliated. Genome sequencing of non-piliated and super-piliated Pph identified few mutations that separate these genotypes from wild type Pph – and none present in genes known to be directly involved in type IV pilus expression. Expression analysis revealed that 81.1% of GO terms up-regulated in the non-piliated strain were down-regulated in the super-piliated strain. This differential expression is particularly prevalent in genes associated with respiration — specifically genes in the tricarboxylic acid cycle (TCA) cycle, aerobic respiration, and acetyl-CoA metabolism. The expression patterns of the TCA pathway appear to be generally up and down-regulated, in non-piliated and super-piliated Pph respectively. As pilus retraction is mediated by an ATP motor, loss of retraction ability might lead to a lower energy draw on the bacterial cell, leading to a different energy balance than wild type. The lower metabolic rate of the super-piliated strain is potentially a result of its loss of ability to retract.

## Introduction

Halo blight is an economically significant disease of leguminous agricultural crops caused by the Gram-negative bacterial pathogen *Pseudomonas syringae* pv. *phaseolicola* (Pph) (Romantschuk and Bamford 1986). The genome of Pph comprises a 5.93 Mb chromosome and two plasmids, 132 kb and 52 kb in size (Joardar *et al*. 2005). Virulence in Pph requires a pathogenicity island located on the larger plasmid (Jackson *et al*. 1999). However, the ability for Pph and other Pseudomonads to attach to plants is via chromosomally-encoded type IV protein pili, which allow the bacterial cells to adhere to external structures (e.g., leaf surfaces) and to resist environmental perturbations such as rain splatter and wind, which could disrupt bacterial invasion of plant tissues (Romantschuk *et al*. 1993; Roine *et al*. 1996; Burdman *et al*. 2011). One of the key mechanisms enabling motility of plant-pathogenic Pseudomonads — fundamental to their interactions with host plants — occurs through twitching motility (see review by Mattick 2002), in which type IV pili are repeatedly extended and retracted to facilitate movement, analogous to grappling hooks. Spontaneous mutants of Pph that lack type IV pili show greatly reduced adherence to leaf surfaces and lower incidence of halo blight disease, whereas virulence of wildtype and pilus-deficient Pph mutants is equivalent when these strains are injected directly into bean leaves (Romantschuk and Bamford 1986). Therefore, adherence ability via type IV pili can be considered a conditional virulence factor directly involved in epiphytic colonization by *P*. *syringae* pathovars such as Pph.

Because type IV pili in Pseudomonads are critical for host plant attachment, it is unsurprising that some bacteriophages have evolved to exploit these conserved structures to facilitate viral infection of bacterial cells (Romantschuk and Bamford 1985). In particular, phage ϕ6 and other members of the dsRNA virus family *Cystoviridae* undergo virulent (lytic) infection of Pph, using the type IV pilus as the initial site of cellular attachment (Roine *et al*. 1996; see review by Mindich 2006). Cystoviruses are characterized as having three dsRNA strands (segments) per particle and a lipid envelope. Attachment of phage ϕ6 to the type IV pili of Pph is mediated by the viral attachment protein P3 (Gottlieb *et al*. 1988; Duffy *et al*. 2006). Once attached, it is believed that retraction of the pilus moves one or more ϕ6 particles in close proximity to the bacterial cell membrane, allowing membrane fusion (Bamford *et al*. 1987) and ultimately phage entry into the cell via the viral lytic enzyme P5 (Gottlieb *et al*. 1988; Dessau *et al*. 2012). Phage ϕ6 then uses the host cell’s metabolism to cause a typical lytic infection, where cell lysis liberates mature virions capable of infecting host cells expressing the type IV pilus (Bamford *et al*. 1976) (see review by Mindich 2006). In addition to the popularity of Pph as a model for studying generalized plant-pathogen interactions (Arnold *et al*. 2011), the infection dynamic between Pph and phage ϕ6 can also be readily examined *in vitro,* facilitating studies of the evolution of host-parasite interactions (Turner and Chao 1999; Lythgoe and Chao 2003; Dennehy, Abedon, *et al*. 2007).

Despite the growing popularity of Pph and phage ϕ6 as a study system in evolutionary biology (Dennehy 2009), the evolution of Pph resistance to phage ϕ6 remains poorly characterized. Laboratory culture of Pph in the presence of phage ϕ6 can be used to select for spontaneous bacterial mutants that are partially or fully resistant to phage infection (Romantschuk and Bamford 1985; Lythgoe and Chao 2003). We are unaware of any study that has characterized the full spectrum of mechanisms that allow Pph resistance to phage ϕ6. However, prior *in vitro* experiments show that evolved resistance generally coincides with changes in the number of type IV pili produced by the bacteria (Romantschuk and Bamford 1985; Lythgoe and Chao 2003), suggesting that type IV pili alteration may be the primary resistance mechanism. Notably, two of these resistant forms differ markedly in pili number: strains lacking type IV pili altogether, and strains with super-abundant type IV pili (Romantschuk and Bamford 1985; Lythgoe and Chao 2003). Resistance in non-piliated strains is due to the elimination of the initial attachment site for phage ϕ6, although some low-level infection of these hosts occurs, presumably due to virus particles colliding directly with the cell surface (Dennehy, Abedon, *et al*. 2007). Super-piliated host strains have elevated numbers of pili compared to wild type Pph, which suggests that these bacteria should be even more sensitive to phage ϕ6 infection. Instead, super-piliated bacteria adsorb relatively larger numbers of phage particles as expected, but remain uninfected apparently because they do not retract their pili (Romantschuk and Bamford 1985; Lythgoe and Chao 2003); this inability to retract type IV pili appears similar to the mechanism described in *P. aeruginosa* (Johnson and Lory 1987). Thus, super-piliated Pph strains ‘soak up’ virus particles in culture, but restrict phage infection because the non-retractable pili prevent virus particles from encountering the cell surface (Lythgoe and Chao 2003; Dennehy, Abedon, *et al*. 2007). Wild-type, non-piliated and super-piliated Pph strains show similar maximal growth rates, stationary-phase densities and relative fitness abilities (i.e., competition for limited nutrients) in liquid culture in the laboratory (Dennehy, Friedenberg, *et al*. 2007). More importantly, super-piliated mutants of Pph are similar to wild-type bacteria in terms of leaf adherence, and initiation of halo blight disease (Romantschuk and Bamford 1986) This observation suggests that super-piliated phage-resistant mutants should be selectively favored over non-piliated (plant-adherence deficient) resistant mutants in the wild; however, we are unaware of any field studies that have directly compared the success of super-piliated versus non-piliated resistance mutants in natural populations of Pph, when exposed to phage attack.

To investigate differences between phage ϕ6 resistant Pph strains and wild-type bacteria, we compared gene-expression profiles and whole-genome sequences of the two major types of resistant strains and wild-type bacteria. In particular, we examined genomic and gene expression variation between ordinary (wild-type) Pph and two spontaneous mutants of Pph that differed in the number of type IV pili expressed per cell: non-piliated, and super-piliated.

## Materials and Methods

### Strains

Wild-type (WT) Pph was obtained from American Type Culture Collection (ATCC strain #21781, a.k.a. strain HB10Y). Non-piliated (P–) Pph strain #LM2509 is a spontaneous mutant of WT Pph, kindly provided by L. Mindich (Public Health Research Institute, Newark, NJ, USA). Super-piliated (P+) Pph strain #KL6 is also a spontaneous mutant of WT Pph, kindly provided by L. Chao (University of California, San Diego, La Jolla, CA).

### Culture conditions

All strains were grown, diluted and plated at 25°C in Luria broth (LB; 10 g NaCl, 10 g Bacto^®^ tryptone [Becton, Dickinson and Co., Sparks, MD], and 5 g Bacto^®^ yeast extract per L), as described in Dennehy *et al*. (Dennehy, Abedon, *et al*. 2007). Cultures were initiated by placing a single colony grown on LB agar into a flask with 10 mL of LB medium. Flasks were shaken at 120 rpm with incubation for 24 hrs, which was sufficient for bacteria to achieve stationary-phase density (∼1.65 × 10^9^ cells/ml).

### Genome sequencing and analysis

Sequencing followed the methods described in O’Brien *et al*. (O’Brien *et al*. 2011). Briefly, DNA was isolated from 1 mL of stationary-phase culture using a Puregene Genomic DNA Purification Kit (Qiagen Canada, Toronto, ON) with the Gram-negative bacterial culture protocol with double volumes of each reagent, repeating the protein precipitation step twice, and spooling the DNA during the precipitation step. DNA was sheared to 200 base pairs (bp) using a Covaris S-series sample preparation system and paired-end sequencing libraries were prepared using sample preparation kits from Illumina (San Diego, CA). Libraries were multiplexed and run on a single lane of an Illumina GA IIx sequencer for 80 cycles per paired-end. Reads were mapped to the genome sequence of wild-type Pph strain HB10Y (Guttman unpublished data) using the CLC Genomics Workbench (Århus, Denmark), and polymorphisms were detected with the CLC Probabilistic Variant Caller. Annotations were copied from the fully sequenced reference strain Pph 1448A (Joardar *et al*. 2005) using a Mauve whole genome alignment (Darling *et al*. 2004).

We evaluated mutational changes in four genes known to be directly involved in type IV pilus formation – type IV pilus assembly protein (PilM), type IV pilus biogenesis protein (PilN), type IV pilus biogenesis protein (PilO) and type IV pilus biogenesis protein (PilP) (Roine *et al*. 1996, 1998; Taguchi and Ichinose 2011; Nguyen *et al*. 2012) in P+ and P- compared to wild type in order to asses any direct genotype – phenotype explanations for the observed phenotypic characteristics of these two strains.

For a well-characterized *P. syringae* pathovar distantly related to Pph (*P. syringae pv. tomato strain* #DC3000), putative sigma70 binding sites inferred from the RegulonDB database (Huerta *et al*. 1998) were downloaded from http://www.ccg.unam.mx/Computational_Genomics/PromoterTools and homologous positions in Pph HB10Y were determined from a whole genome alignment made with Mauve (Darling *et al*. 2004). Mutations occurring within 50bp upstream of the inferred transcription start sites were identified using BEDTools (Quinlan and Hall 2010). Pph HB10Y homologs were identified for 5121 out of 6716 putative sigma70 binding sites in *P. tomato* DC3000, and mutations were present within 50 bp of the putative transcription start sites for 8 (7 in P+ strain KL6, 1 in P– strain LM2509).

### Gene expression analysis

Two biological replicates of each strain were cultured. For RNA extraction we used a standard Trizol approach. The bacterial cells (roughly 10^7^ to 10^8^ cells per ml of Trizol) were lysed in Trizol (Life Technology) and homogenized with a micropestle (Eppendorf), and RNAs were suspended in DEPC treated water. RNA quality was checked via spectrophotometry (NanoDrop). Reverse transcription of the RNA into cDNA was performed using the Invitrogen SuperScript Double-Stranded cDNA Synthesis kit (Thermo Fisher) using random primers. For each biological replicate, a technical replicate of the reverse transcription was performed using multi-targeted primers (Adomas *et al*. 2010) designed for the Pph genome, instead of random primers, for first strand synthesis. All cDNA samples were labeled using a NimbleGen One-Color (Cy3) DNA Labeling Kit (Madison, WI, USA). Hybridization of cDNA to NimbleGen 4 x 72k arrays (090326_P_syringe_B728a_expr; NimbleGen design ID #9510; Madison, WI, USA) according to the manufacturer guidelines was conducted by the Yale Center for Genome Analysis (New Haven, Connecticut, USA). Microarray slides were scanned with a GenePix 4000B (Axon Instruments, Foster City, CA). Spots were located and expression was quantified following NimbleGen Arrays User’s Guide, and spots with unusual morphology or with erratic signal intensity distribution were excluded from all analyses. Raw fluorescence data were normalized as in Townsend (Townsend 2004), and normalized data were then statistically analyzed using Bayesian Analysis of Gene Expression Levels (BAGEL, (Townsend and Hartl 2002; Townsend 2004)). Genes were considered expressed and deemed well measured only when the median spot foreground exceeded the median background plus two standard deviations of the background intensity. Well measured genes were considered as significantly differentially expressed when P ≤ 0.05. Significantly up and down regulated genes with Gene Ontology (GO) terms were used to determine significantly enriched GO terms using AmiGO 1.8 (Carbon *et al*. 2009) with a significance level of 0.01 and a minimum of two genes for each enriched term. The JCVI CMR (Davidsen *et al*. 2009) database was used for background filtering and results were visualized using the AmiGO GO graph tool (Carbon *et al*. 2009). Visualization of gene expression differences among strains was done by using the clustergram function of the bioinformatics toolbox of the MatLab computational mathematics suite (Mathworks, Natick, MA, USA). The data were first mean-normalized and then transformed so that gene expression values were represented as standard deviations above or below the mean. Data were not sorted for statistical significance beforehand although later conclusions did take specific *P* values between strains into account.

## Results

### Strain isolation

The super-piliated and non-piliated mutants in the current study are spontaneous mutants of WT Pph strain HB10Y, obtained via laboratory assays that isolated spontaneous resistance mutants in bacterial populations subjected to phage ϕ6 attack. This design facilitated genomic and gene-expression comparisons among strains, which may have been complicated if the strains were more distantly related, such as drawn from disparate natural sources and/or geographic locations. We noted that all three strains achieved similar stationary-phase densities (∼1.65 × 10^9^ cells/ml) and maximal growth rates in Luria broth (Dennehy *et al*. 2007), indicating that average number of type IV pili expressed per cell constituted the major differences between WT Pph and each of the two spontaneous mutants.

### Genomics of super-piliated and non-piliated bacterial strains

Illumina-based genome sequencing using 80 base pair (bp) paired-end reads of the super-piliated (P+) and non-piliated (P–) strains produced 1.9 and 3.6 million reads respectively. Subsequent referenced alignment to the wildtype (WT) *P. syringae pv. phaseolicola* strain HB10Y resulted in a mean coverage of 14.0× and 24.4× for P+ and P–.

Alignment of the P+ strain identified 97 mutations separating it from WT (Table 1, Table S1), consisting of 26 large-scale deletions, five deletion-insertion polymorphisms (DIPs), seven multi-nucleotide variants (MNVs) and 59 single nucleotide polymorphisms (SNPs). A notably large deletion of 1039 bp occurred in a retrotransposon hot spot (rhs) family gene, with the remainder averaging 120bp (54–269 bp). Fourteen of the deletions occurred in inter-genic regions, six in characterized genes and six in hypothetical proteins of unknown function. All five DIPs in P+ occurred in intergenic regions; three DIPs were in consecutive positions 158 bp downstream of the gene encoding the type III effector HopQ1. The remaining two DIPs were upstream of a prophage and fructose-1, 6-bisphosphate aldolase gene. Each observed MNV consisted of a 2–3bp change, six of which occurred in intergenic regions and the remaining MNV located in the *tolQ2* ORF, causing a threonine to leucine amino-acid substitution at the 121^st^ position. Of the 59 SNPs detected in P+, 15 occurred in ORFs, of which six were non-synonymous amino acid changes.

**Table 1:** A summary of genomic and gene expression differences between super-piliated and non-piliated Pph.

| Mutation Type | Strain |
| --- | --- |

|  | P+ | P– |
| --- | --- | --- |
| Large Scale deletions | 26 | 4 |
| Deletion-Insertion |  |  |
| Polymorphisms | 5 | 8 |
| Multi-nucleotide Variants | 7 | 8 |
| Single Nucleotide |  |  |
| Polymorphisms | 59 | 62 |
| Total | 97 | 82 |
| Expression Differences |  |  |
| Genes UP–regulated | 503 | 104 |
| Genes Down-regulated | 451 | 28 |
| Gene Ontology Terms UP–regulated | 4 | 3 |
| Gene Ontology Terms Down-regulated | 38 | 1 |

Alignment of the P– strain identified 82 total mutations separating it from WT (Table 1, Table S2) consisting of four large-scale deletions, eight DIPs, eight MNVs, and 62 SNPs. A single large deletion of 2116 bp occurred in the gene encoding the *rulA* protein, with the remainder averaging 80 bp (53–112 bp). Four of the eight observed DIPs occurred in a GntR family transcriptional regulator, causing frameshift mutations at amino acids 131 and 132. Two DIPs occurred in a putative H+ antiporter subunit C causing a frameshift at Phe_50_, and one DIP was present in the *PilX* ORF causing a frameshift mutation at Ile_192_. The remaining DIP was present in an intergenic region. The eight observed MNVs ranged from 2–5bp in length. Five of the eight MNVs were in intergenic regions. However, the remaining three MNVs were located within the H+ antiporter, a GntR family transcriptional regulator and a TolQ protein causing frameshifts at Phe_50_, at Asp_131_ and an amino acid change from Thr_121_ to Leu, respectively. Of the 62 SNPs observed in P–, 34 were located in intergenic regions, and of the 28 SNPs occurring in confirmed ORFs, 16 represented non-synonymous changes.

None of the observed mutations were present in any of the genes known to be directly involved in type IV pilus formation (i.e., PilM, PilN, PilO and PilP) (Roine *et al*. 1996, 1998; Taguchi and Ichinose 2011; Nguyen *et al*. 2012) in P+ or P-.

Putative phage ϕ6 binding site mutations are shown in Table 2. One was detected in P–, with seven detected in P+. The single binding site mutation in P– represents a SNP (G to C) upstream of a citrate transporter gene, and downstream of a hypothetical protein. The binding site mutations in P+ include an MNV upstream of type III effector *HopQ2* which was also significantly upregulated in comparison with WT, along with six deletions ranging in size from 64–269 bp. Of the genes potentially affected by these deletions, a *LuxR* family D--binding response regulator and an iron(III) dicitrate transport protein fecA were significantly downregulated in P+, and a ribosomal subunit interface protein, a periplasmic amino acid-binding protein, and a hypothetical protein were significantly upregulated in P+.

**Table 2:**
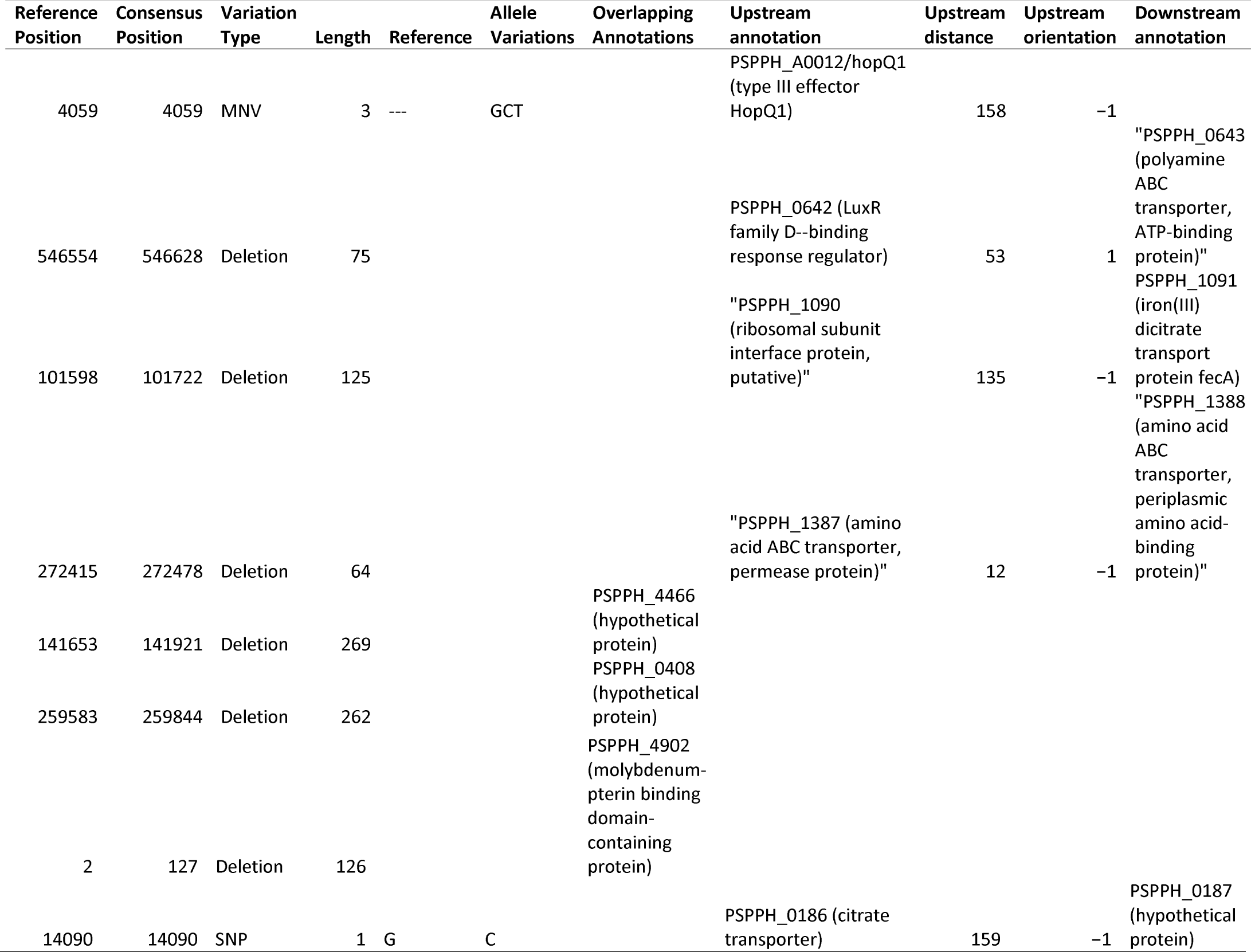
Details putative sigma70 binding sites inferred from the RegulonDB database for which significant regulatory changes were detec ted.

### Transcriptome Analysis

The gene expression (GE) profiles of the 3 strains were compared using a clustergram and heatmap (Figure 1) analysis applied to all genes (N = 5069), and applied to genes that showed a significant difference (P < 0.05) between strains (N = 1009). Both analyses showed that the GE profile of P+ differed more substantially from WT than did P– (Figure 1) with higher significance when restricted to significantly different genes. Two major clusters of genes were observed: one cluster contained genes highly expressed in P+ compared to WT and P–, and another contained genes highly expressed in WT and P– in comparison with P+.

**Figure 1:**
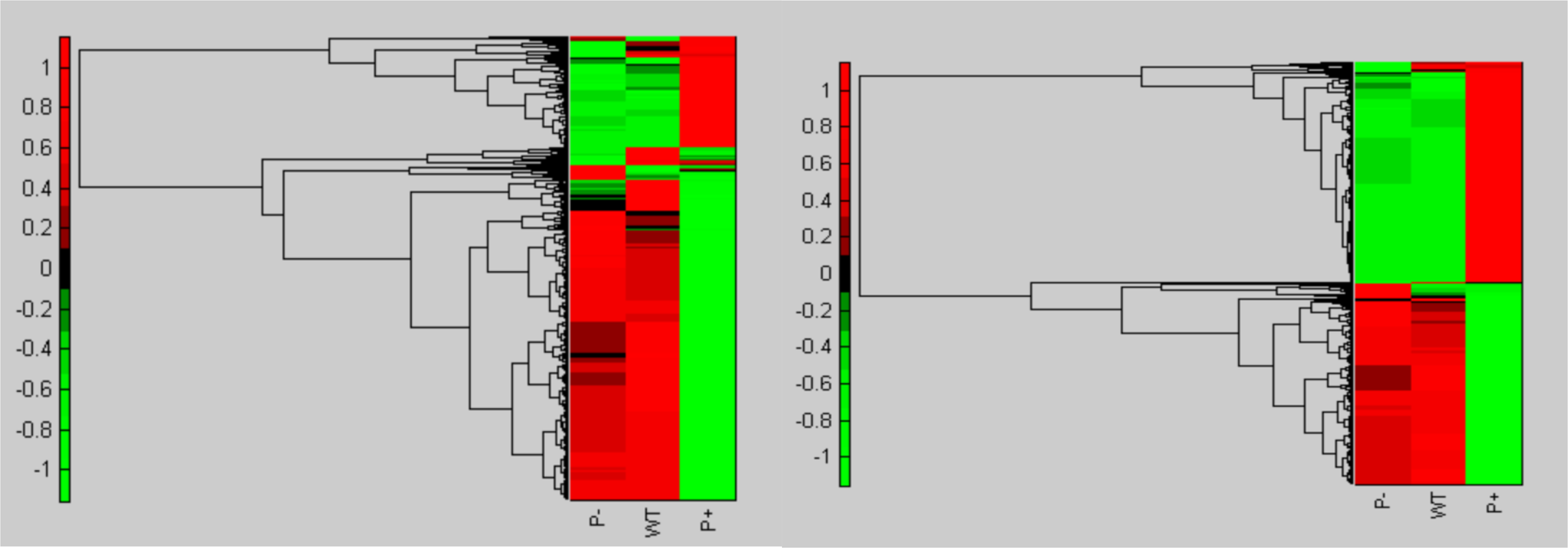
Clustergram of standardized values of all genes (left) and significantly (P < 0.05) differentially expressed genes (right). Significantly differentially expressed genes are those that are significantly up-regulated (red) /down-regulated (green) in P+ or P– relative to WT.

Of the 5069 genes measured on the microarray, 954 genes (18.8%) showed significant expression-level differences between WT and P+, with 503 genes upregulated and 451 genes downregulated (Figure 2, Table 1, Table S3). To further examine genome-wide differences in GE, genes upregulated in P+ were analyzed for GO term enrichment (Figure S1). We observed that 10 terms were significantly enriched, all of which pertained to processes of transposition and DNA metabolism. An enrichment analysis of genes down-regulated in P+ showed significant enrichment for 48 different GO terms. Visualizing the organization of these GO terms revealed that a few key biological processes were enriched: gene expression and translation, nucleotide and ribonucleotide biosynthesis, and cellular respiration (Figure S2). Of the 5069 genes measured on the microarray, 132 (2.6%) genes significantly differed in gene expression levels between WT and P–. Of these 132 genes, 104 genes were up-regulated and 28 genes were down-regulated (Table S1). We found that 49 GO terms were enriched for P– up-regulation and 31 GO terms were enriched for down-regulation. Visualization of these GO terms showed particularly high enrichment for intracellular organelles and ribosomal structure in genes up-regulated in P– (Figure S3), and genes associated with conjugation as well as protein transport and secretion showed high levels of down-regulation in P– (Figure S4).

**Figure 2:**
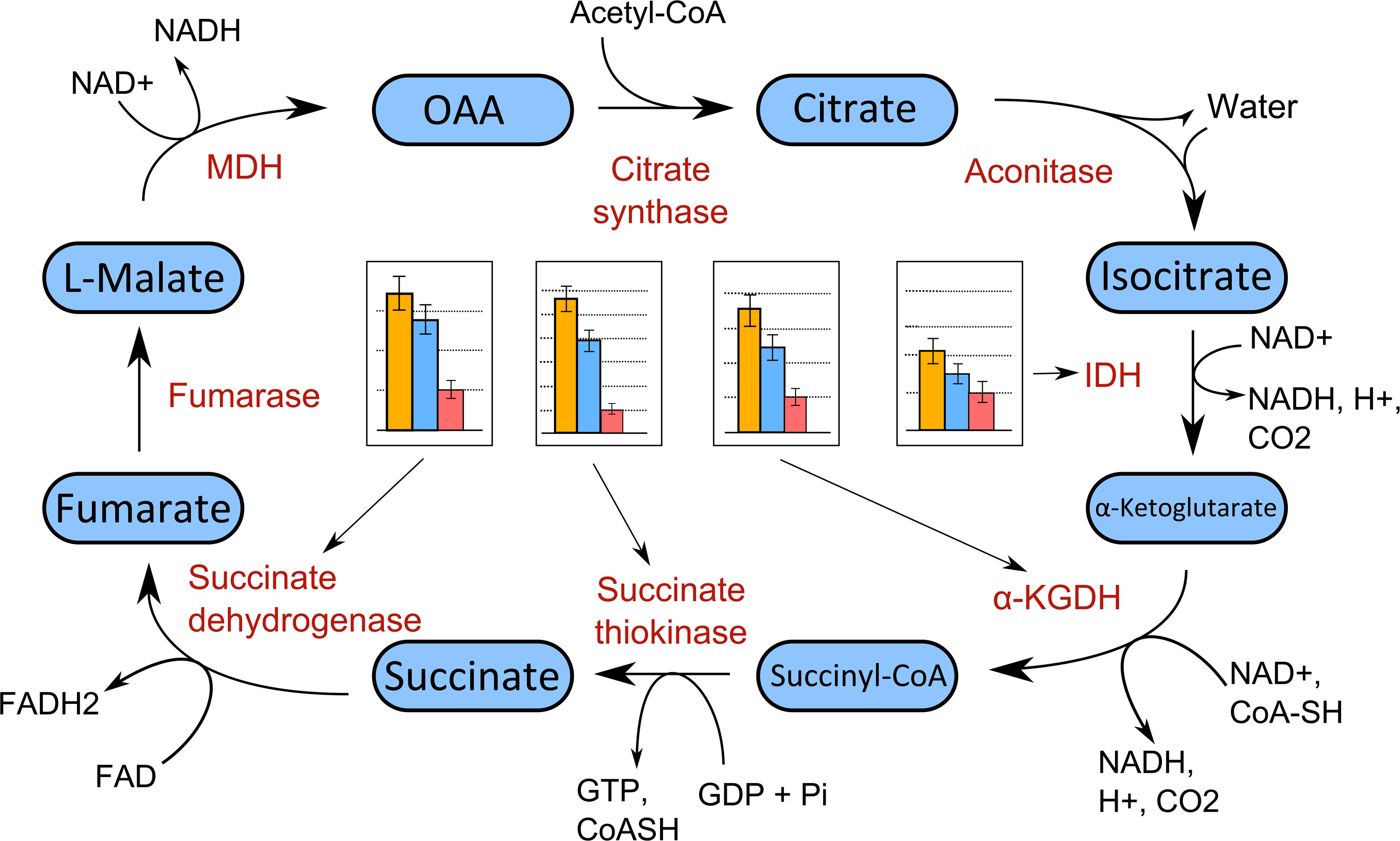
Expression levels of key enzymes of the TCA cycle in P– (yellow), WT (blue), and P+ (red) strains.

## Discussion

### Genomic differences between P+ and P– phage resistant P. syringae pv. phaseolicola

Genome sequencing of P+ and P– strains identified 97 and 82 mutations that differed from WT strain, respectively. These mutations consisted of 26 deletions, five DIPs, seven MNVs and 59 single SNPs in P+, and four deletions, eight DIPs, eight MNVs, and 62 SNPs in P– (Table 1). Of these differences, 20 and 27 mutations resulted in non-synonymous changes to protein coding genes in P+ and P– strains. These observations constitute a relatively very small number of protein coding changes — approximately one mutation per 300 kb and 220 kb respectively, indicating that the observed differences in pilus number associated with P+ and P– stem from a small number of large effect mutational changes. Genes affected by protein coding changes in P+ included SNPs located in a general virulence surface antigen protein, and a type III effector AvrB4-2—both loci are associated with plant-disease virulence in Pph (Arnold *et al*. 2011). Genes affected by protein coding changes in P– were a large deletion in the RulA protein, which confers UV resistance in Pph (Sundin *et al*. 2000). Both P+ and P– had MNVs in the *tolQ2* gene— including identical mutations from threonine to leucine at the 121^st^ position. *tolQRA* genes encode membrane proteins that determine phage sensitivity versus resistance in *Escherichia col*i bacteria, where *tolQRA* mutants were resistant to phage infection (Picken and Beacham 1977). It is therefore possible that mutations in TolQ membrane proteins play a role in both entry of phage ϕ6 into Pph cells, as well as resistance of Pph to phage ϕ6 attack. The specifics of TolQ mutations associated with phage resistance deserve further investigation. Interestingly, we did not find any mutations in genes known to be required for pilus expression (i.e. PilM, PilN, PilO and PilP) (Roine *et al*. 1996, 1998; Taguchi and Ichinose 2011; Nguyen *et al*. 2012). This indicates that the link between the P+ and P- phenotypes and their corresponding genotypes is likely to be complex and possibly regulatory in nature.

### Interactions between mutation and gene expression

Of the 20 non-synonymous mutations in P+, six were located in genes with significant expression differences compared to WT; four were up-regulated and two were down-regulated. Of the four genes which were up-regulated, two were annotated as hypothetical proteins of unknown function. However, the two remaining upregulated genes were related to Pph virulence. The first is AvrB4-2, a type III effector protein associated with Pph virulence in common bean (*Phaseolus vulgaris*) plants (Zumaquero *et al*. 2010). P+ had a SNP mutation in AvrB4-2 that caused an amino acid change from aspartic acid to asparagine at the 240^th^ position. *Avr* genes in *P. syringae* are involved in host-specific resistance mechanisms between the bacterium and its plant hosts (Lee *et al*. 2004). The second SNP was initially identified in a search of 19 *P. syringae* genome sequences for genes related to bacterial pathogenicity (Baltrus *et al*. 2011). A valine at the third amino acid position of the gene was changed to a leucine.

In P–, only one of the 26 genes containing non-synonymous mutations showed a significant expression difference compared to WT: a phage integrase-family recombinase. In this gene, two mutations were observed: a change from leucine to proline at position 216, and a change from methionine to leucine at position 219. Also of interest is the combination of a binding site mutation and up-regulation of a type III effector *hopQ2* gene in P–. Type III virulence effector proteins are important for *P. syringae* to infect and grow in plant hosts, and thus these mutations may yield insights into the virulence of P– strains of *P. syringae* (i.e., aside from the expected reduced conditional virulence on plant hosts, due to the lack of type IV pili in P– strains).

### *Gene expression differences between P+ and P– phage resistant P. syringae pv.* phaseolicola

GE Profiling demonstrated that P+ had significantly more genes both up and down-regulated in comparison to WT than P–: compared to WT, 503 and 104 genes were significantly up-regulated and 451 and 28 genes were downregulated in P+ and P–. GO term enrichment analysis revealed four and 40 GO terms enriched for up-regulation in P+ and P–, and 38 and one GO terms enriched for down-regulation in P+ and P–.

The exceptionally low number of GO terms enriched for up-regulation in P+ suggests that transcription and DNA metabolism are likely to be key processes in P+ resistance to phage infection, and the genes annotated for each term represent candidates for further, gene specific analysis. Down-regulation of genes associated with gene expression and translation, nucleotide and ribonucleotide biosynthesis, and cellular respiration in P+ (Figure 2) indicates the potential importance of these pathways in P+ resistance to phage infection.

The up-regulation of genes involved in gene expression and ribosome production in P– suggests the up-regulation of protein production in general. Additionally, the up-regulation of genes associated with cellular respiration and ATP synthesis indicates that energy production in general increased in P–. The observed down-regulation of pili-related genes in P– is unsurprising given that this strain lacks pili, however the specific functions of the proteins within this down-regulated set of genes have yet to be investigated.

Perhaps of most interest is the observation that 81.1% of GO terms up-regulated in P– were shown to be down-regulated in P+ (Figure 1, Table S3). This result indicates that regulatory changes in similar pathways have led to divergent, independent mechanisms of phage resistance in Pph. A number of genes associated with respiration – specifically TCA cycle, aerobic respiration and acetyl-CoA metabolism genes – were up-regulated in P– and downregulated in P+.

There were 10 and seven genes annotated for all three of these GO terms differentially expressed in comparison with WT in P+ and P– respectively, of which four were common to both. The four genes were succinate dehydrogenase, alpha-ketoglutarate dehydrogenase, isocitrate dehydrogenase, and succinyl-CoA synthetase – all encoding key enzymes in the TCA cycle. Plotting the expression levels of these genes reveals a similar pattern of greatest expression in P– and lowest expression in P+ (Figure 2). Viewing these genes in the context of their location in the TCA cycle (Figure 2) reveals that the enzymes they encode are responsible for catalyzing energy-yielding reactions, either in terms of a nucleoside triphosphate or a reduced cofactor. In general, kinetic and allosteric feedback loops between metabolites and enzymes in a biological pathway generally maintain a degree of stability, and the increases or reductions in the activity of one gene versus another does not significantly change flux through the pathway. However, if all of the enzymes are up-regulated/down-regulated in concert, such as in the case of P+ and P–, the whole pathway would be affected. Thus, the expression pattern of these four genes suggests that the TCA pathway is generally up-regulated/down-regulated in P+ and P– respectively, and that its pathway flux relates to pilus regulation

The down-regulation of the TCA pathway could indicate lower metabolic rate of the P+ strain, potentially a result of its loss of retraction ability. Pilus retraction is mediated by an ATP motor, so loss of retraction ability could lead to a draw of ATP on the bacterial system (Helaine *et al*. 2007). These conclusions are speculative, however, and require further experiments for confirmation.

The divergent expression patterns of these TCA cycle genes in P– versus P+ reflect the general reciprocal GE profiles of the two strains. Overall, genes up-regulated in P– are down-regulated in P+ and vice versa (Table S3). The only exception to these reciprocal GE profiles is that transposition is up-regulated in both P– and P+. The fact that in P– the transposition genes are up-regulated, while pili genes are down-regulated may suggest a mutation causing a fundamental disconnect between these two otherwise co-regulated processes. This observation will be a continued area of study, since the exact regulatory mechanisms behind transposition in Pph are not well known.

The results of our study are useful for general research efforts harnessing phage ϕ6 and Pph. In particular, these microbial strains have been used in prior studies in experimental ecology and evolutionary biology, especially to conduct experimental evolution that addressed fundamental questions in these fields (Turner and Chao 1999; Montville *et al*. 2005; Dessau *et al*. 2012; Goldhill and Turner 2014). However, these prior studies have not focused on co-evolutionary interactions between phage ϕ6 and Pph, perhaps because resulting genetic changes in the host bacteria could not be compared to existing full genome sequences of the founding strains. Our study addresses this limitation and lends more power to the phage ϕ6/Pph model system, because such work can now harness affordable re-sequencing approaches to compare ancestral and derived strains of bacteria. Also, recent studies show that Cystoviruses are readily found in the terrestrial phyllosphere in temperate environments, especially in association with bacteria residing on or within common legumes such as white clover plants (Mindich *et al*. 1999; Silander *et al*. 2005; O’Keefe *et al*. 2010; Díaz-Muñoz *et al*. 2013). These studies emphasize the prevalence of RNA phages in terrestrial biomes (Mindich *et al*. 1999; Silander *et al*. 2005; O’Keefe *et al*. 2010; Díaz-Muñoz *et al*. 2013), and to our knowledge Cystoviruses constitute the only non-marine RNA phages that are actively researched in the wild (Culley *et al*. 2006). Whereas these earlier studies were used to infer population structure and isolation-by-distance in Cystoviruses, our study is useful for future empirical work that manipulates Cystoviruses and Pph strains on legumes in field experiments or in the greenhouse. For example, the bacteria characterized in our study could be used to examine the effects of phage infection on the community dynamics of bacteria that differ in plant pathogenicity, under realistic conditions such as on leaf surfaces. Overall, characterization of Pph pili mutants in the current study support the crucial need to further develop emerging models in phage/bacteria interactions, which should broaden the understanding of symbiotic interactions between the most prevalent entities in the biosphere (Hendrix 2002; Wasik and Turner 2013).

## Acknowledgements

We thank members of the Turner and Townsend labs for providing valuable feedback on the study. This work was supported by grant #1021243 to P.E.T. from the U.S. National Science Foundation. D.P. was supported by the Beckman Scholars Program of the Arnold and Mabel Beckman Foundation.

## Supplementary Material

**Table S1:** Genomic mutations present in super-piliated Pph in comparison to WT.

**Table S2:** Genomic mutations present in non-piliated Pph in comparison to WT.

**Table S3:** Gene expression differences between super and non-piliated Pph in comparison to WT. The number 1 indicates genes which are significantly up or down regulated, 0 indicates genes which do not show a significant difference.

**Figure S1:**
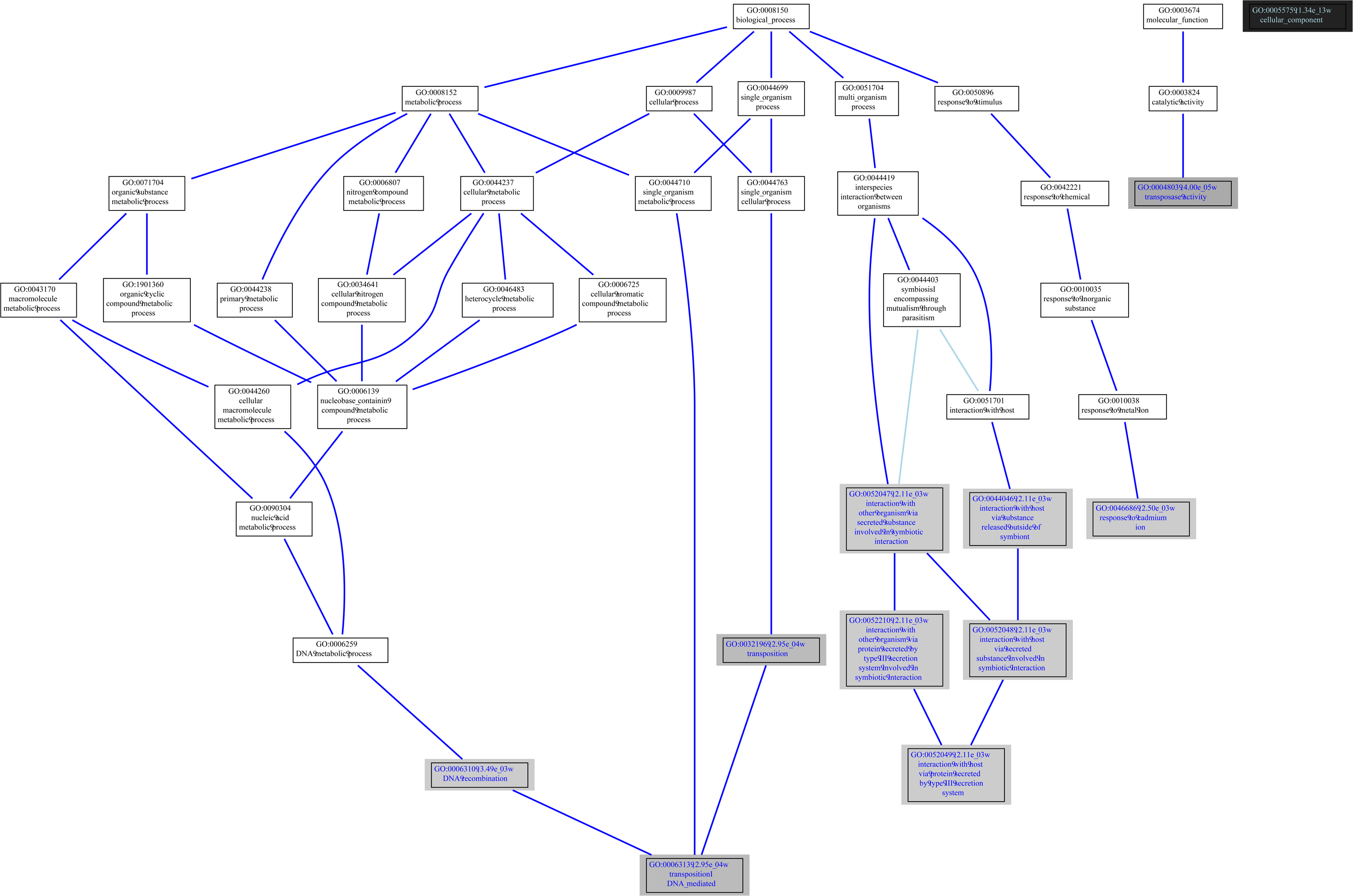
Visualization of the GO terms enriched in the genes up-regulated in P+. Shading corresponds to level of significance. The vertical axis generally indicates greater or lesser specificity of the GO term because of the hierarchical definitions of the terms. The three major biological functions that are enriched are those of Gene expression and translation, nucleotide and ribonucelotide biosynthesis, and respiration. The majority of unboxed terms are mostly general terms lacking a unifying biological function.

**Figure S2:**
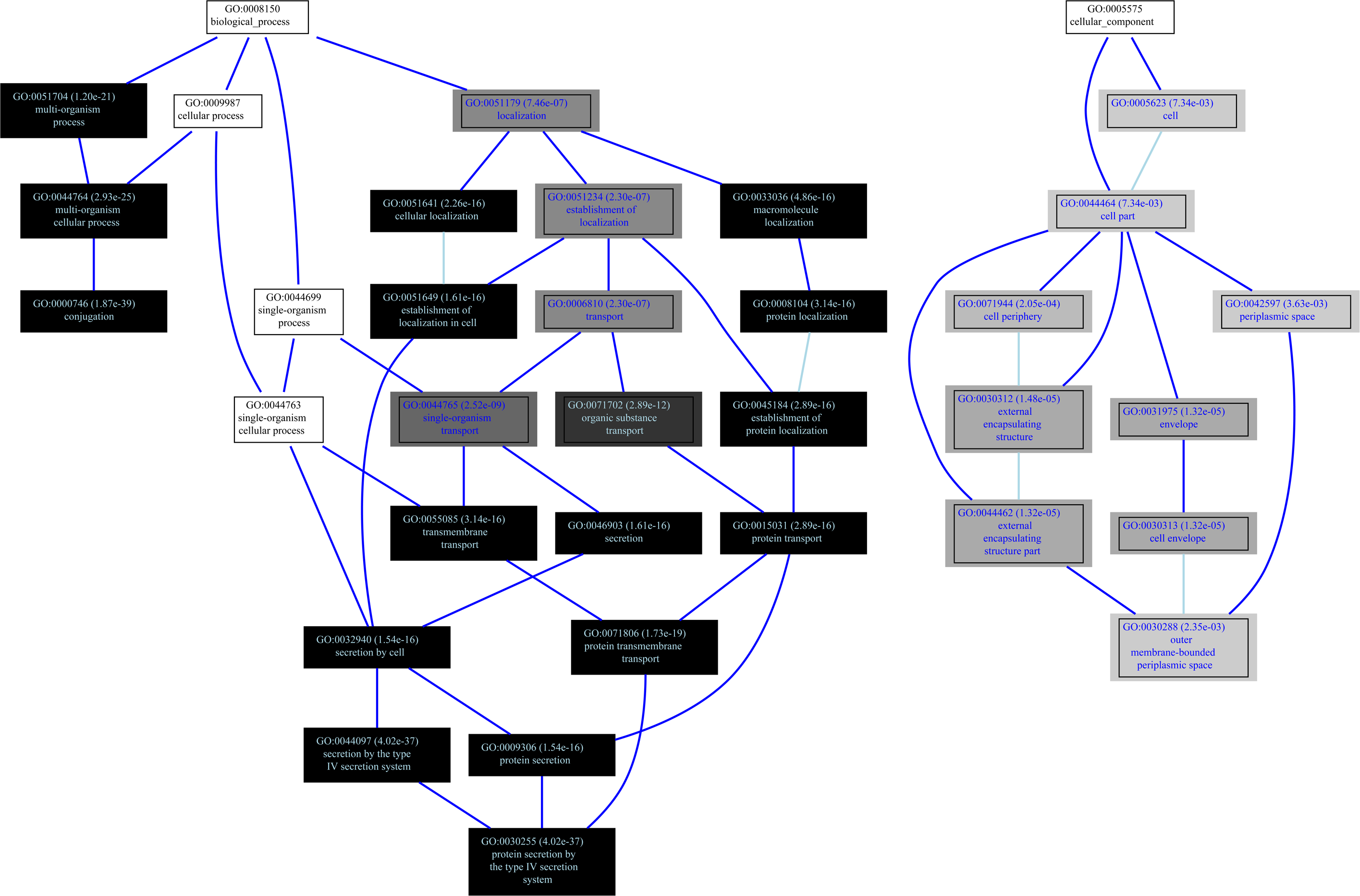
Visualization of the GO terms enriched in the genes down-regulated in P+. Shading corresponds to level of significance. The vertical axis generally indicates greater or lesser specificity of the GO term because of the hierarchical definitions of the terms. The three major biological functions that are enriched are those of gene expression and translation, nucleotide and ribonucelotide biosynthesis, and respiration. The majority of unboxed terms are mostly general terms lacking a unifying biological function.

**Figure S3:**
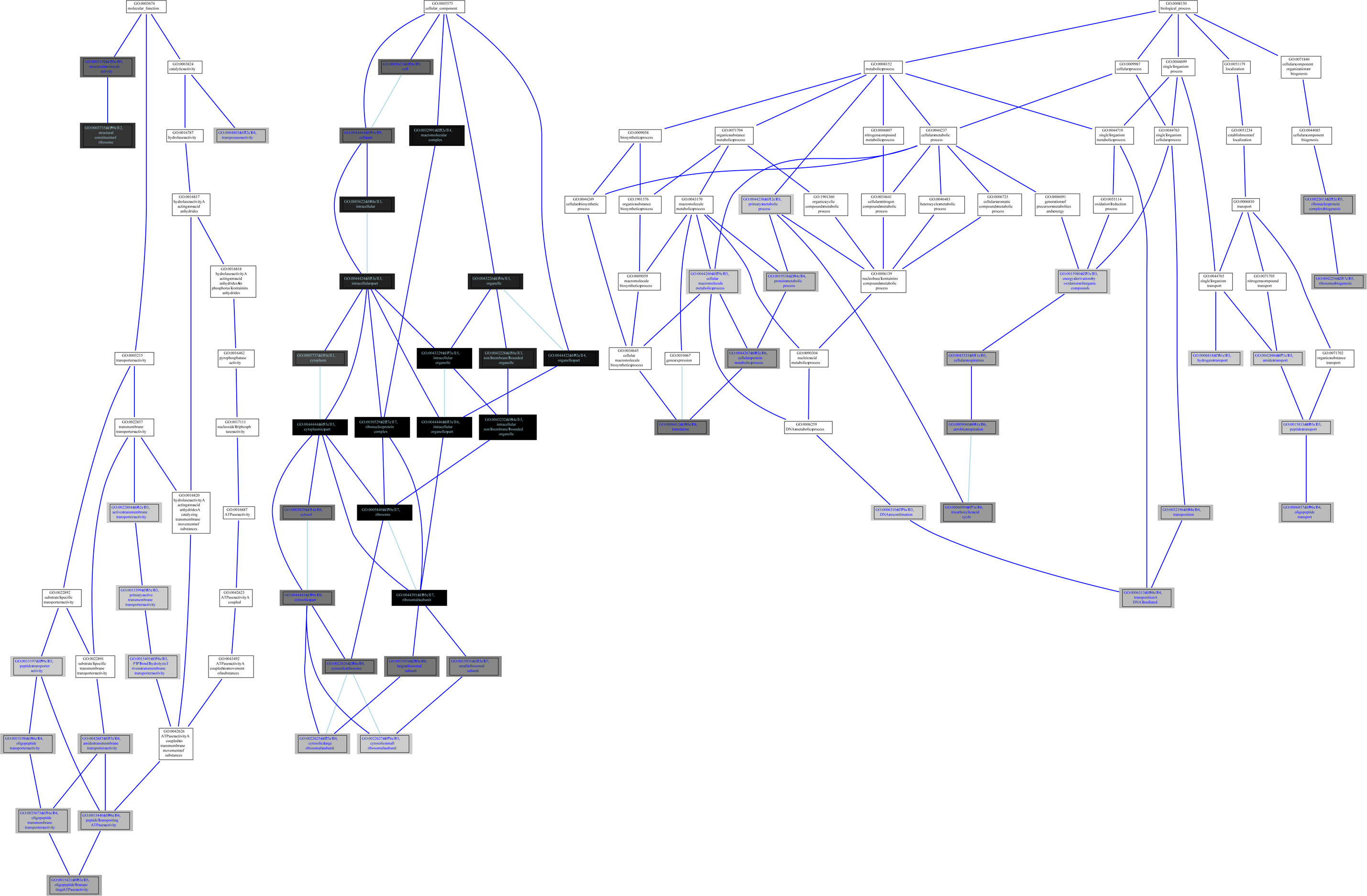
Visualization of the GO terms enriched in the genes upregulated in P–. Shading corresponds to level of significance. The vertical axis generally indicates greater or lesser specificity of the GO term because of the hierarchical definitions of the terms. The three major biological functions that are enriched are those of gene expression and translation, nucleotide and ribonucelotide biosynthesis, and respiration. The majority of unboxed terms are mostly general terms lacking a unifying biological function.

**Figure S4:**
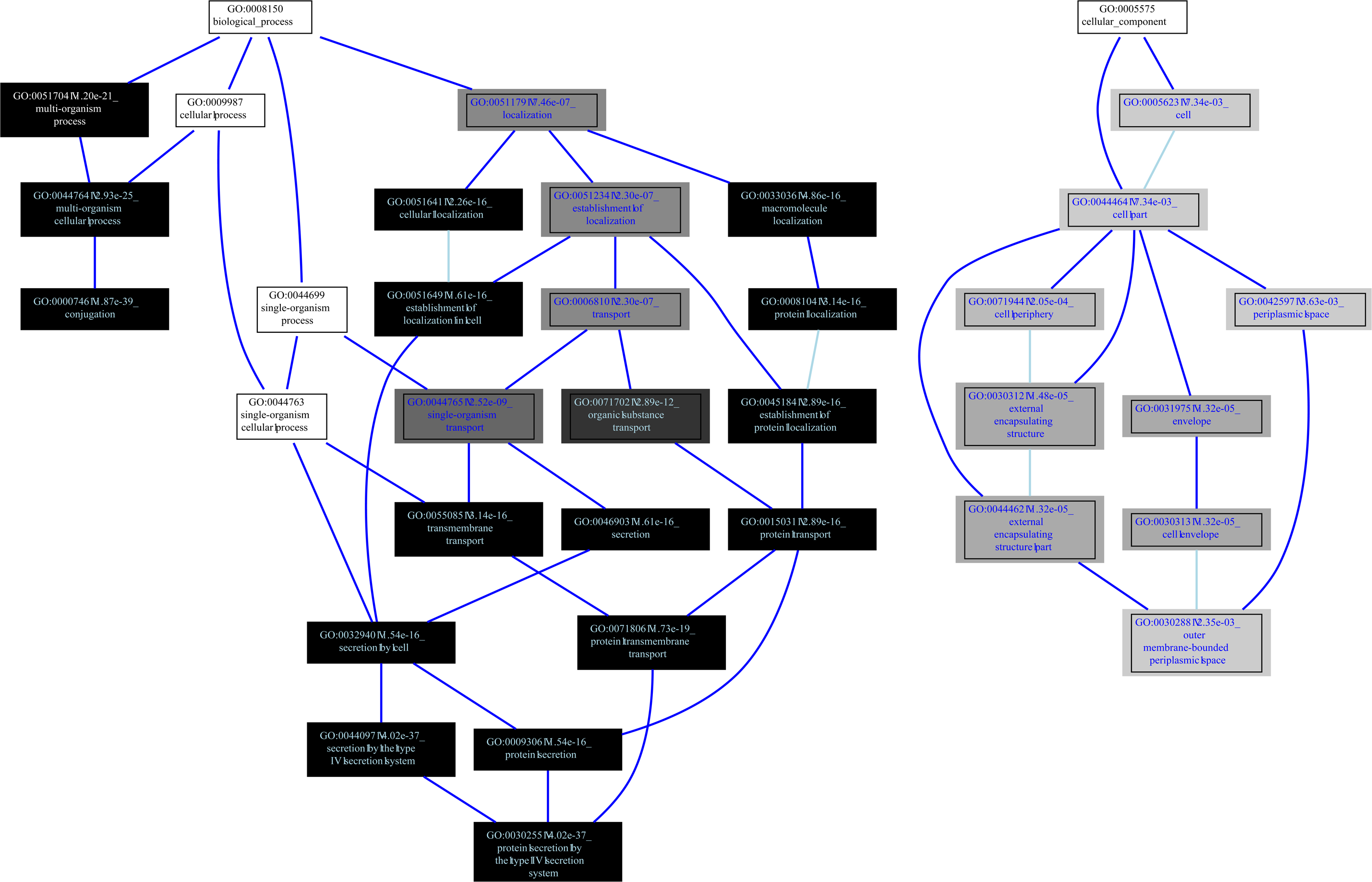
Visualization of the GO terms enriched in the genes downregulated in P–. Shading corresponds to level of significance. The vertical axis generally indicates greater or lesser specificity of the GO term because of the hierarchical definitions of the terms. The three major biological functions that are enriched are those of gene expression and translation, nucleotide and ribonucelotide biosynthesis, and respiration. The majority of unboxed terms are mostly general terms lacking a unifying biological function.

